# ATOM-1: A Foundation Model for RNA Structure and Function Built on Chemical Mapping Data

**DOI:** 10.1101/2023.12.13.571579

**Authors:** Nicholas Boyd, Brandon M. Anderson, Brent Townshend, Ryan Chow, Connor J. Stephens, Ramya Rangan, Matias Kaplan, Meredith Corley, Akshay Tambe, Yuzu Ido, Jake Yukich, Tabitha Tcheau, Ayah Abdeldayem, Gabriel Ferns, Harsh Patel, Shaon Barman, April Schleck, Adrian L. Sanborn, Stephan Eismann, Raphael J. L. Townshend

## Abstract

RNA-based medicines and RNA-targeting drugs are emerging as promising new approaches for treating disease. Optimizing these therapeutics by naive experimental screening is a time-consuming and expensive process, while rational design requires an accurate understanding of the structure and function of RNA. To address this design challenge, we present ATOM-1, the first RNA foundation model trained on chemical mapping data, enabled by data collection strategies purposely developed for machine learning training. Using small probe neural networks on top of ATOM-1 embeddings, we demonstrate that this model has developed rich internal representations of RNA. Trained on limited amounts of additional data, these small networks achieve state-of-the-art accuracy on key RNA prediction tasks, suggesting that this approach can enable the design of therapies across the RNA landscape.

## 1 Introduction

RNA-based medicines have recently demonstrated significant therapeutic potential through the successful development of mRNA vaccines, antisense oligonucleotides (ASOs), siRNAs, and RNA editing therapies [1–6]. In addition, small molecule drugs targeting endogenous RNA species offer new avenues to treat disease, in particular when the corresponding protein targets are undruggable [7, 8]. Realizing the full therapeutic potential of RNA requires predicting and optimizing complex properties, whether the stability of mRNA vaccines, activity of ASOs, or the binding affinity of small molecules to RNA. As many of these properties are mediated by structure and experimental structure determination is difficult, computational models that understand structure are important to accelerate the development of RNA-focused therapies.

A major challenge in the design of RNA-focused therapies is the lack of ground truth data to use for modeling. Functional data, such as on siRNA toxicity, can often only be collected at low throughput. With respect to structural data, few experimentally determined tertiary structures of RNA are available. In fact, only 1% of entries in the Protein Data Bank (PDB) comprise RNA alone [9], despite the over 10-fold excess of genome intervals that produce RNA relative to proteins [10]. While evolutionary information encoded in multiple sequence alignments (MSAs) can provide critical insights on structure and function, these alignments are often shallow and uninformative for human targets and engineered sequences [11]. Consequently, state-of-the-art RNA structure and function prediction approaches fall short of the recent successes of highly accurate protein prediction methods [12].

Transfer learning from foundation models pretrained on large datasets has proven successful in many data-limited applications of machine learning [13]. Key to success on downstream tasks is the emergence of complex internal representations that encode a general understanding of the application domain. One technique to demonstrate the emergence of these internal representations is the use of so-called *probe networks*: small neural networks that take as input internal embeddings from the larger model and produce predictions of properties of interest [14–17]. A foundation model with a rich and accessible internal representation of RNA structure could enable accurate predictions even for severely data-limited tasks. Critically, training such a model requires a sufficiently large and informative dataset.

We show that *chemical mapping* can provide such a dataset for an RNA foundation model. In a chemical mapping experiment, chemical reagents can be used to modify RNA nucleotides in a structure-dependent manner [18–21]. These modifications are detected by sequencing to glean information on an RNA’s conformational states in solution or in cells. Multiplexing over RNA species combined with next-generation sequencing (NGS) allow for the collection of large datasets. Additionally, and in contrast to MSAs, chemical mapping experiments can be run on arbitrary RNA sequences, allowing the exploration of sequence space beyond natural sequences. As these experiments directly measure structural information, foundation models trained on chemical mapping data could enable better predictions on structure-related tasks for RNAs of interest compared to models trained on natural sequences alone [22–24].

We present ATOM-1, a foundation model trained on large quantities of chemical mapping data collected in-house across different experimental conditions, chemical reagents, and sequence libraries. Using probe networks, we show that ATOM-1 has developed rich and accessible internal representations of RNA. Despite their size, these small probe networks demonstrate state-of-the-art accuracy on several tasks, including predicting RNA 3D structure, secondary structure, and in-solution RNA stability.

## 2 Results

### 2.1 Training a foundation model on chemical mapping data

We first give a brief overview of chemical mapping and how to pose it as a supervised machine learning problem. Chemical mapping experiments modify RNA and produce a collection of sequencing reads for each input RNA species; each read may contain substitutions, insertions, or deletions relative to the original sequence (Figure 1). The distribution of these mutations is related to the structure (or ensemble of structures) of the input RNA; different chemical mapping reagents and experimental conditions measure different aspects of RNA structure. For many of these reagents, a first-order approximation is that unpaired nucleotides are more likely to result in mutations than paired nucleotides.

**Figure 1:**
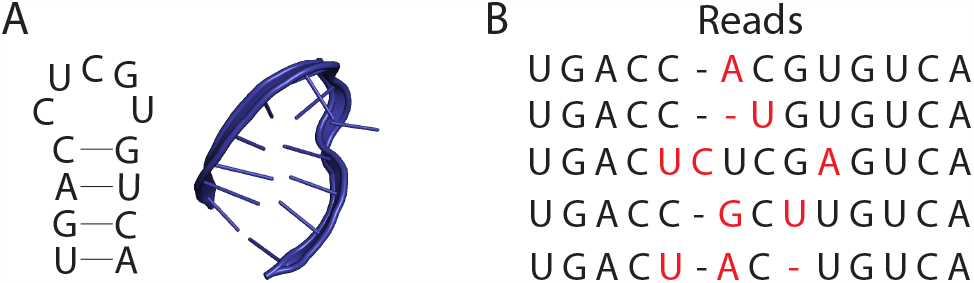
RNA structure and chemical mapping reads. **(A)** An RNA secondary structure (left) and tertiary structure (right) for the same RNA. Lines in the secondary structure denote base-paired positions. Many chemical mapping reagents will preferentially, but not exclusively, modify unpaired positions. **(B)** Sequencing reads from a chemical mapping experiment, with mutations (red) from the original sequence occurring more frequently at unpaired positions.

From a machine learning perspective this is a standard sequence-to-sequence problem: the input sequence is the RNA species, while the output sequences are the observed reads assigned to that species. Readout via NGS allows the input species to be multiplexed and experiments to be scaled to produce the hundreds of billions of tokens needed to train high-capacity machine learning models.

We collect chemical mapping data using several chemical reagents on a set of diverse, custom-designed libraries under several different conditions. To this data we fit a custom, structure-aware, encoder-decoder sequence-to-sequence transformer-based model. For an RNA sequence of length *n*, the embedding produced by the encoder is two objects: the *single representation*, which is an array of size *n*-by-512, and the *pair representation*, an array of size *n*-by-*n*-by-256 [25]. In the following sections we show that the encoder’s embeddings contain rich and accessible information on RNA structure and function.

In machine learning, probe networks are commonly used to demonstrate the emergence of accurate and accessible representations in large, pretrained models [26]. Importantly, computational probing experiments emulate the process of prototyping the use of the foundation model for a new prediction task. A typical probing experiment consists of two steps. First, we train a small network (the probe) to predict the property of interest directly from the foundation model embeddings. Next, to show that performance of the probe is the direct result of the foundation model and not our training procedure or probe network, we train the same network without access to embeddings (the baseline). If the performance of the probe is substantially better than that of the baseline we conclude that the foundation model contains useful and accessible representations of the property of interest.

### 2.2 Secondary structure prediction

RNA secondary structure is characterized by patterns of hydrogen bonding between nucleotide bases in canonical Watson-Crick or wobble base pairs [29]. These structures are crucial for RNAs’ biological function and the design of RNA-focused therapies. From a mathematical standpoint, a secondary structure *S* of an RNA of length *n* is a set of unordered pairs *{i, j}* where *i≠ j ∈* 1, …, *n*. Each pair in *S* is called a base pair.

To demonstrate that ATOM-1 has an understanding of secondary structure, we consider probe networks that take embeddings from ATOM-1 as input. Since base pairing is a property of each pair of nucleotides, it is natural to apply these probes to the pair representation independently along the last dimension. As an example, a 257-parameter linear model trained on *a single secondary structure* yields qualitatively-reasonable predictions of secondary structure (Figure 2). In fact, despite only being trained on an FMN riboswitch aptamer structure (PDB ID: 6WJR [28], 112 nucleotides), this simple probe is able to generalize to distinct RNA classes, for instance a cloverleaf-like RNA domain (PDB ID: 8DP3 [27], 90 nucleotides). This demonstrates that in the process of learning to predict chemical mapping data, ATOM-1 has developed an accessible representation of secondary structure.

**Figure 2:**
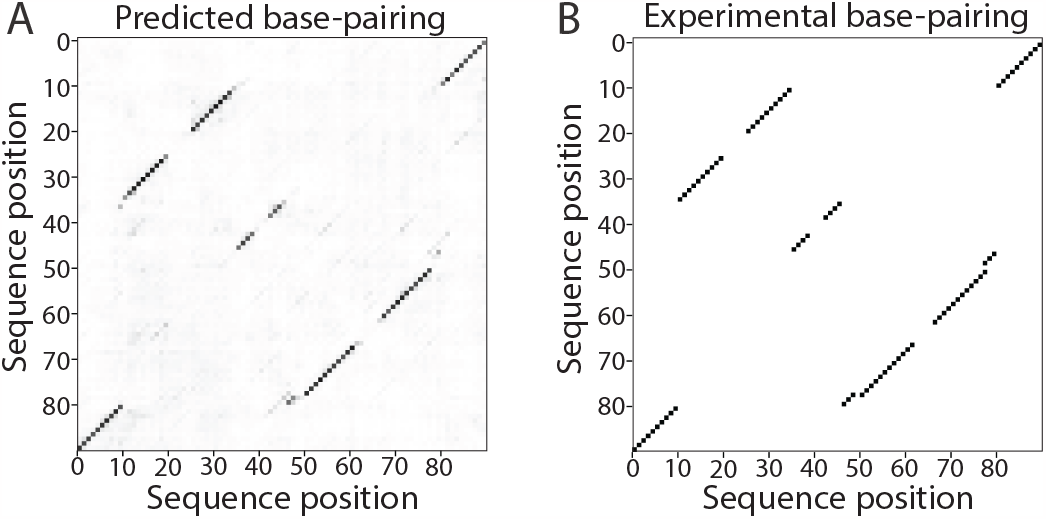
A linear probe with 257 parameters trained on one secondary structure generalizes to other RNAs. (**A**) The predicted probability of each base pair for PDB ID 8DP3 [27] as estimated by our 257-parameter probe. (**B**) The ground truth secondary structure for PDB ID 8DP3 represented as a symmetric matrix of base pairs. This linear probe of ATOM-1’s pair representation was trained on a single secondary structure (PDB ID: 6WJR [28]). The accurate prediction demonstrates that ATOM-1 has developed accessible and accurate representations of secondary structure.

To show that the secondary structure representations developed by ATOM-1 are highly accurate, we consider a slightly more expressive probe: a multilayer perceptron (MLP) with a single hidden layer of dimension 2048 (for a total of *∼*2.6M parameters). For comparison, we consider a probe with the same architecture applied to RNA-FM [22], a foundation model trained on naturally-occurring RNA sequences; following section 2.1, we include a baseline network with the same architecture applied only to sequence features. For technical details see section S1.

We train the probe networks on a subset of single-chain RNA secondary structures derived from PDB entries before April 30, 2020. For testing, we use secondary structures from PDB entries published after May 1, 2020 and further exclude sequences with more than 80% sequence identity to our training set from the evaluation. See section S1.2 for more details. Figure 3A presents the accuracies of the different prediction methods as measured by F1-score (see section S1.1). The probe of ATOM-1 is competitive with physics-inspired methods, RNAFold [33] and CONTRAFold [34], and performs substantially better than the same probe architecture applied to RNA-FM. Our baseline—the probe architecture applied directly to sequence features—demonstrates minimal prediction accuracy.

**Figure 3:**
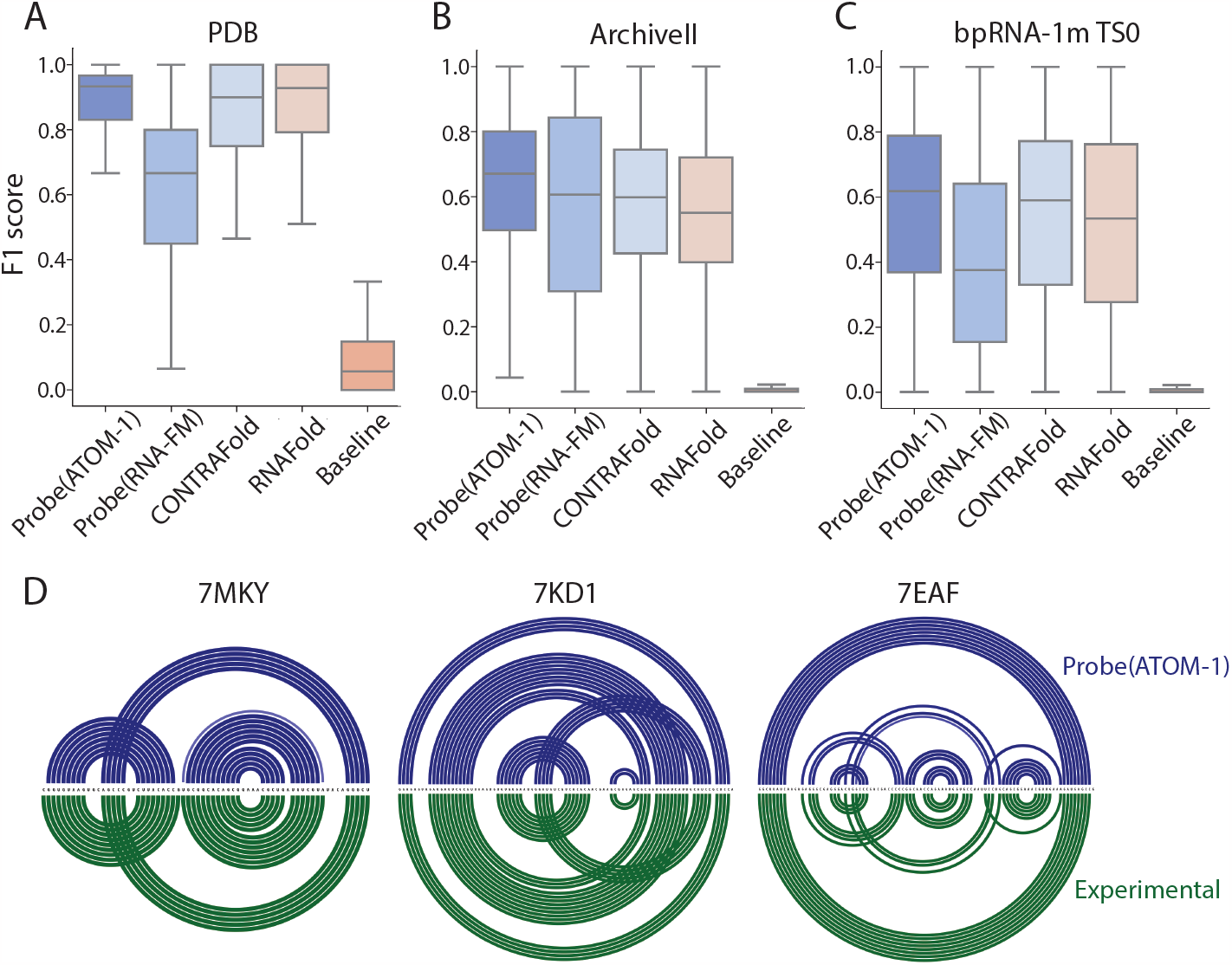
A probe of ATOM-1 for secondary structure prediction generalizes with high accuracy. We train single-hidden-layer MLP probe networks on embeddings from ATOM-1 and RNA-FM. As a baseline, we also train the same architecture without access to any foundation model as a baseline. We further include comparisons with two physics-inspired predictors, CONTRAFold and RNAFold, that do not use foundation models. Comparisons are shown for evaluation sets derived from three sources: **(A)** the PDB, **(B)** Archive II, and **(C)** bpRNA-1m TS0. For all three panels, the probe networks were trained on PDB structures. Unlike for the PDB evaluation set, the secondary structures in ArchiveII and bpRNA-1m TS0 are inferred from multiple sequence alignments. **(D)** Arc diagrams comparing secondary structures predicted using the probe of ATOM-1 to experimental secondary structures derived from the PDB. Comparisons are shown for three structures from the PDB evaluation set: the SARS-CoV-2 frameshift stimulation element (PDB ID: 7MKY [30]), an apo THR riboswitch aptamer (PDB ID: 7KD1 [31]), and a SAM-I riboswitch variant (PDB ID: 7EAF [32]). Arcs connect nucleotides in Watson-Crick base pairs. The intensity of coloring represents the predicted probability of base-pairing.

To test the generalization capability of our probe, we validate on two additional datasets: bpRNA1m TS0 [35, 36] and ArchiveII [37]. As with the PDB evaluation set, we remove test cases with high sequence identity to our training set. Secondary structure in these datasets is not derived from experimentally-determined tertiary structure, but inferred from multiple-sequence alignments. Despite the shift in domain, our model remains highly accurate, demonstrating strong generalization ability (Figure 3B,C).

We find that our probe generates accurate predictions for complex RNAs across diverse RNA classes and lengths (Figure 3D). For instance, we accurately predict secondary structures for a SARS-CoV-2 frameshift stimulation element construct, an apo THR riboswitch aptamer, and a SAM-I riboswitch variant. These examples demonstrate that the probe is able to correctly predict pseudoknots, secondary structure elements which physics-inspired methods often fail to predict [33, 34].

Finally, we note that our probe technique is purely *local*: each prediction for a pair of residues uses only the single and pairwise representation for those two residues. This is in contrast to previous secondary structure techniques which use non-local dynamic programming algorithms [33, 34], repeated convolutional layers with large receptive fields [22, 35, 40], or both [41, 42]. Because our probe network does not include any interactions between nucleotides, any predictive performance originates from the representation present in the ATOM-1 embeddings alone.

### 2.3 Tertiary structure prediction

While secondary structure is an important aspect of RNA, many therapeutically-relevant properties of RNA are mediated by the full tertiary (3D) structure. A natural question, then, is to what extent ATOM-1 contains readily-accessible 3D structural information, especially since one might suspect that chemical mapping data is dependent only on secondary structure. To answer this, we probe ATOM-1 using a shallow (two-layer), MSA-free variant of the Evoformer [25] with a custom structure module (see section S2). We train and evaluate our model on RNA structures from the PDB and report results on clusters of test set sequences grouped by sequence similarity (see section S2.4).

Figure 4A compares our probe of ATOM-1 to two state-of-the-art 3D structure prediction methods: RhoFold [38], the deep learning method with best performance from CASP15 [43], and RoseTTAFold2NA [39]. Notably, both RhoFold and RoseTTAFold2NA make use of MSAs which are time-consuming to generate and are often unavailable for RNAs of interest [11]. Despite having no access to MSAs and being considerably smaller (*∼*15M parameters) and shallower (2 layers) than RhoFold (*∼*100M parameters in 12 layers) and RoseTTAFold2NA (*∼*68M parameters in 40 layers), our probe produces predictions with higher global accuracy as measured by root mean-squared deviation (RMSD) [44] to experimental structures (Figure 4A). Moreover, compared to our baseline network, which uses an identical architecture without ATOM-1 embeddings, the probe produces predictions with consistently higher local accuracy as measured by the local distance difference test (LDDT) [45] (Figure 4B). Overall, our probe generates the best 3D structure predictions more often than state-of-the-art deep learning methods as measured by both RMSD and LDDT (Figure 4C). Together, these comparisons show that ATOM-1 produces readily accessible and accurate representations of RNA 3D structure.

**Figure 4:**
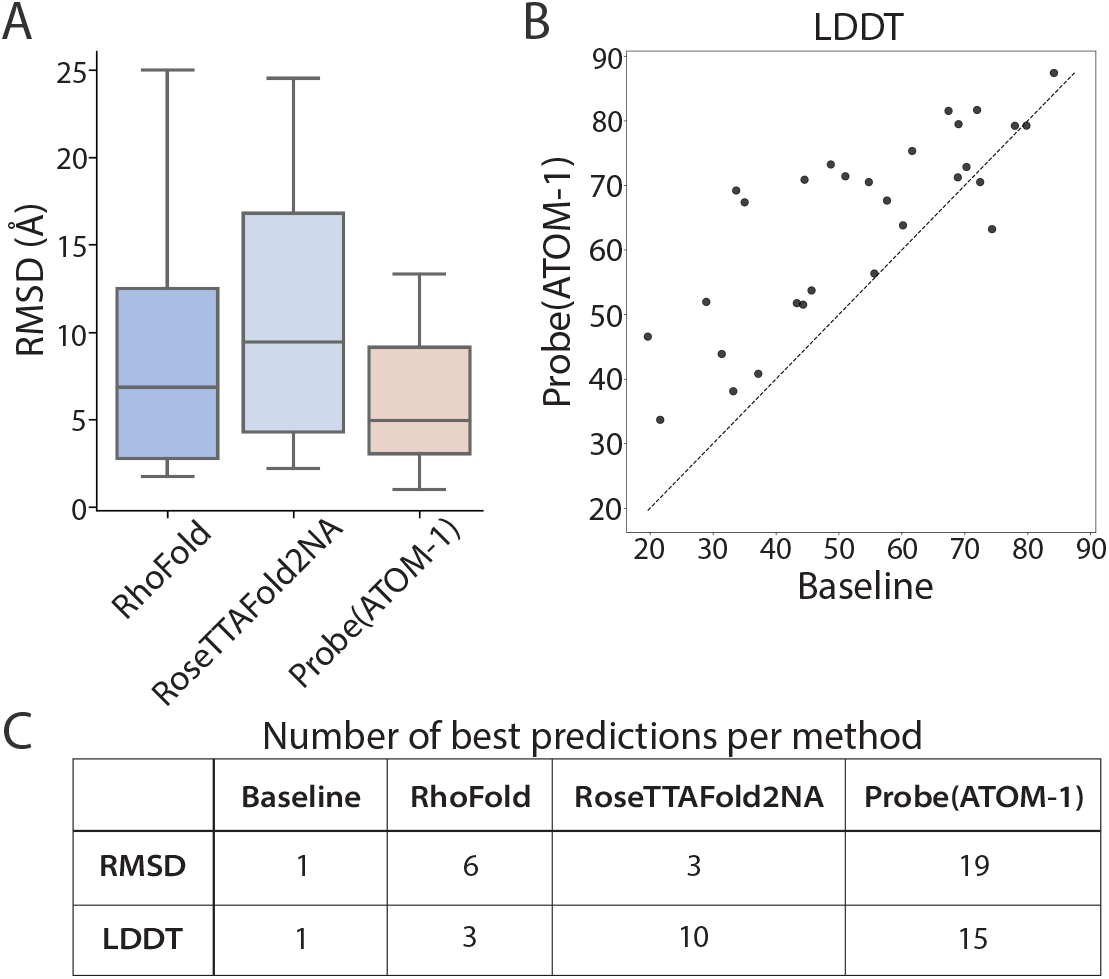
A probe of ATOM-1 for tertiary structure prediction demonstrates state-of-the-art accuracy. Results are shown for *N* = 29 clusters of test set structures published after May 1, 2022. **(A)** Structure prediction accuracy of a probe of ATOM-1 as measured by RMSD compared to RhoFold [38] and RoseTTAFold2NA [39], both of which have access to MSAs. **(B)** Structure prediction accuracy of a probe of ATOM-1 as measured by LDDT versus the baseline model. The baseline is a model with an identical architecture to the probe but without access to ATOM-1. **(C)** Number of cases for which each method predicts the best structure among all tested methods as measured by RMSD and LDDT.

The utility of ATOM-1 embeddings is further evident in the visualizations of predicted 3D structures in Figure 5. We find that our probe network produces RNA models that match the native global fold for diverse RNA targets across a broad range of sequence lengths. These predictions substantially outperform the baseline model without ATOM-1. Notably, this improvement is apparent even in cases where the native structure includes mostly non-canonical base-pairing (for instance, the G-quadruplex in Figure 5B), suggesting that ATOM-1 embeddings contain structural information beyond secondary structure.

**Figure 5:**
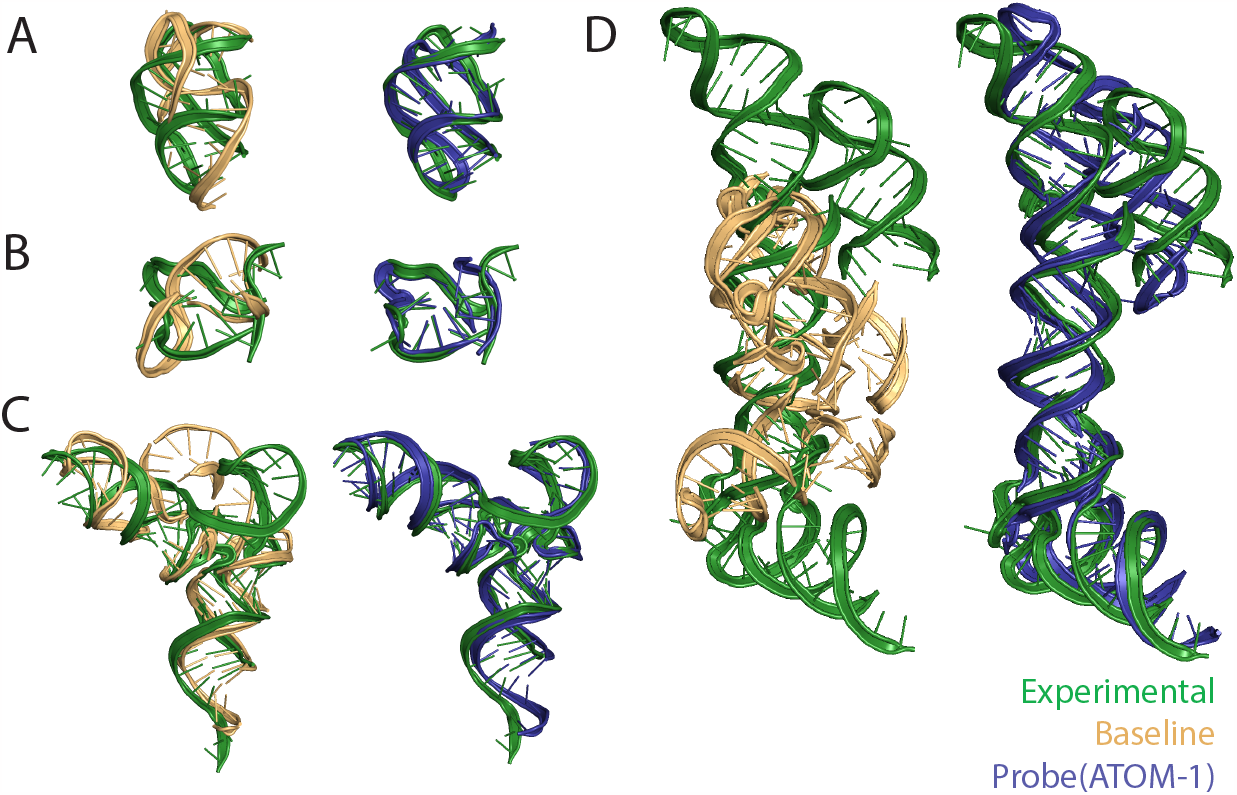
Structure predictions for a probe of ATOM-1 compared to the baseline without foundation model embeddings. The baseline model is identical to our probe architecture but does not use ATOM-1. Predictions are overlaid on experimental structures for different test set RNAs: **(A)** a Pre-Q1 riboswitch (PDB ID: 8FB3 [46]), **(B)** a G-quadruplex (PDB ID: 7SXP [47]), **(C)** a synthetic tRNA (PDB ID: 7URI [48]), and **(D)** a cloverleaf RNA fused with a tRNA (PDB ID: 8S95 [49]).

### 2.4 In-solution stability

Successful distribution of mRNA vaccines requires mRNA constructs that are stable over long periods of time in solution. We evaluate the ability of our foundation model to help predict RNA stability using data from the Stanford OpenVaccine Kaggle community prediction challenge [50].

We train a simple probe network (*∼*10M parameters) to predict degradation and reactivity characteristics from the embeddings of ATOM-1. Figure 6A shows that our method outperforms all 1636 challenge submissions. For comparison, we also show the accuracy of a baseline network without access to ATOM-1 embeddings. As in previous tasks, we observe significant accuracy regression—the test loss of the baseline network is 37% higher compared to the ATOM-1 probe—indicating that the high prediction accuracy of our probe of ATOM-1 is not driven by the probe architecture or training procedure. We provide more details on the prediction task and its evaluation in section S3.2.

**Figure 6:**
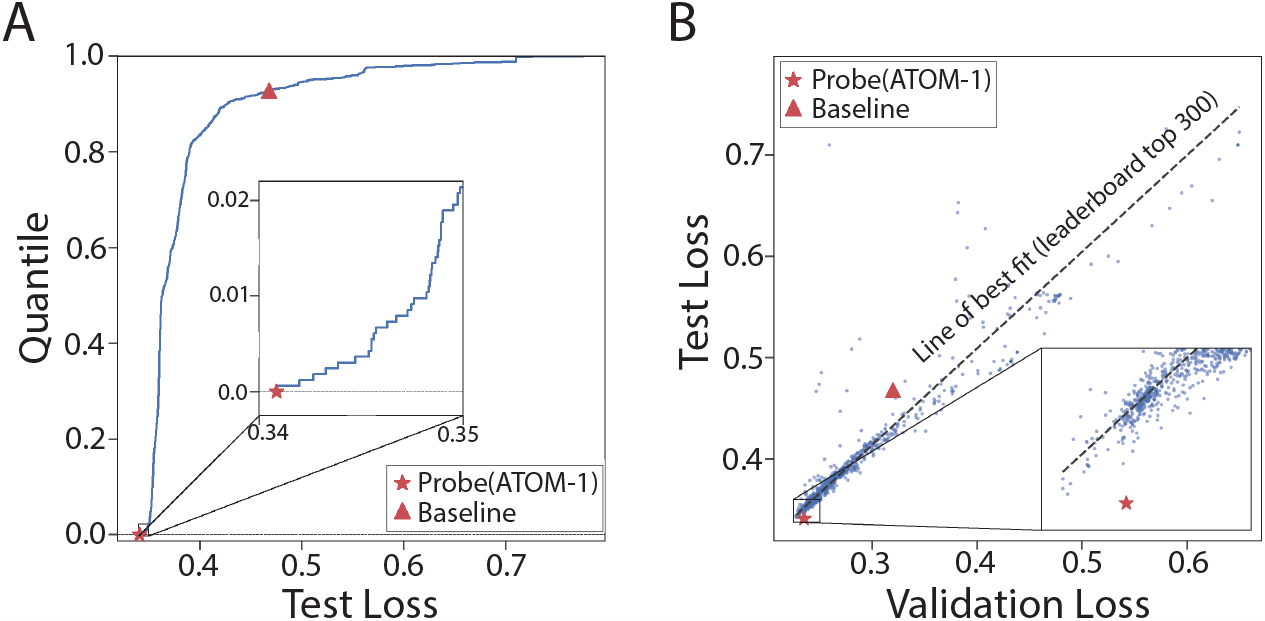
A probe of ATOM-1 takes first place on data from the OpenVaccine community challenge. **(A)** The empirical distribution of test losses across challenge submissions. The quantile value denotes the fraction of submissions with smaller (better) test losses. Lower quantile values indicate better performance. We also show the performance of a baseline with identical architecture to our probe but without access to ATOM-1. **(B)** Validation versus test loss for all submissions from the OpenVaccine Kaggle challenge and our probe model (lower is better). Compared to other methods, we generalize better to long sequences present in the test set. The black dashed line is a line of best fit on the top 300 submissions by test loss. Loss is calculated as the mean prediction RMSE across multiple prediction tasks (see section S3.2).

The design of this challenge allows us to showcase the generalization abilities of models built on top of ATOM-1. Figure 6B compares validation and test losses for the different methods that participated in the challenge. We note that the ATOM-1 probe does particularly well with respect to the sequences in the test set, which are about 30% longer than those in the training and validation sets. During the challenge, participants were able to repeatedly evaluate the accuracy of their methods on the validation set, likely leading to overfitting to this validation set by some methods, whereas an evaluation on the test set was not available until the end of the challenge.

Furthermore, we note that we do not perform any pretraining or self-distillation using test set sequences, whereas the top Kaggle solutions used one or both of these approaches [51, 52]. While these methods are perfectly valid within the confines of the challenge, they are likely to lead to test metrics that are overly optimistic with respect to the prospective performance of models on new sequences—even those drawn from the same distribution as the test set.

## 3 Outlook

We show that ATOM-1, a foundation model trained on large quantities of chemical mapping data, has developed accurate and accessible representations of RNA. Despite their small size, probe networks trained on ATOM-1 embeddings demonstrate state-of-the-art accuracy across multiple tasks. On 3D RNA structure prediction in particular, probing ATOM-1 improves upon methods that have access to coevolution information in multiple sequence alignments—information that is often not available for prospective RNA design or for human RNA targets. On a community challenge for RNA stability prediction, a small ATOM-1 probe takes first place in a retrospective analysis.

ATOM-1’s strong generalization abilities suggest broad applicability across a wide range of other properties relevant to the design of RNA-focused therapies, such as RNA translation efficiency, siRNA toxicity, and ASO activity. Given a small dataset of experimental measurements, the foundation model enables fast and accurate prototyping.

Here we have focused on small probe networks, which are ideal to query the accessibility and information content of foundation model embeddings, but may not yield the highest prediction accuracy. Larger, more expressive networks and more advanced transfer learning techniques can substantially improve this accuracy. Similarly, ATOM-1 is trained solely on data from chemical mapping experiments; this training data can be readily extended to include additional data from experiments that provide orthogonal information on RNA structure and function, enriching the information content of the foundation model embeddings and its generalization abilities.

## Supplementary Material

### S1 Secondary structure probing

For both ATOM-1 and RNA-FM, our probe network consists of a multilayer perceptron with a single hidden layer of dimension 2048. For each pair of nucleotides *i* and *j* we apply the probe to the length 1024 vector *E*_*ij*_ = (*S*_*i*_, *S*_*j*_, *P*_*ij*_). *S*_*i*_ and *P*_*ij*_ are the 512 and 256 dimension single and pair representations produced by ATOM-1. When probing RNA-FM, we used the 640 dimensional representation of the last layer as the single representation. Since RNA-FM does not have a natural pair representation, we concatenated the 20 attention heads from all twelve layers, for a total effective pairwise dimension of 240. As such, the probe network for RNA-FM and ATOM-1 have slightly different numbers of parameters at 3.1M and 2.6M respectively. The output of our probe is a pairing probability *p*_*ij*_ for each pair of nucleotides. We train on the ground truth pairing matrix using the binary cross entropy loss function.

As a baseline, we use a model with the same architecture (a multilayer perceptron with a single hidden layer of dimension 2048) but with sequence features instead of ATOM-1 embeddings as input. The sequence features for pair *ij* are the one-hot embeddings of the nucleotides *i* and *j* and a one-hot embedding of the distance between *i* and *j* in the sequence, *i − j*, with maximum distance of 32 [25]. This model resulted in nearly zero accuracy. To strengthen the baseline in the main text, we applied Sinkhorn’s algorithm to the predicted logits to generate a doubly-stochastic base-pairing probability matrix *p*_*ij*_ such that Σ _*i*_ *p*_*ij*_ = Σ_*j*_ *p*_*ij*_ = 1. The Sinkhorn layer allows the baseline network some non-local interaction between base pairs, improving predictive performance.

#### S1.1 Metrics

To calculate the F1 score for a prediction, we first solve the assignment problem using the Hungarian algorithm to generate the single most probable secondary structure conformation. For this conformation we calculate the Positive Predictive Value (PPV) and Sensitivity (SEN) compared to the ground truth conformation: 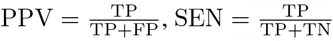 where TP, TN, and FP are respectively the number of true positives, true negatives, and false positives. Finally, the F1 score is the harmonic mean of PPV and SEN : F1 = 2 *×* PPV *×* SEN*/*(PPV + SEN). See Table S1 for a comparison of all three metrics across the three secondary structure test sets.

#### S1.2 Datasets and splits

Our training dataset is a subset of PDB entries described in Sec. S2.4, filtered to include only singlechain RNA structures of unmodified nucleotides. We construct the PDB secondary structure pairing matrices using DSSR [53]. We allow each nucleotide to have only a single pair; if a nucleotide is identified as participating in multiple pairs we use the canonical Watson-Crick pair. For our PDB test set, we use structures published after May 1, 2020. For all three test sets (PDB, ArchiveII, bpRNA-1m TS0), we applied a filter that removed entries with less than 80% sequence identity to our PDB training set. We further cluster each test set at the level of 80% sequence identity, and the resulting F1 metric is reported after averaging over each cluster.

**Table S1:**
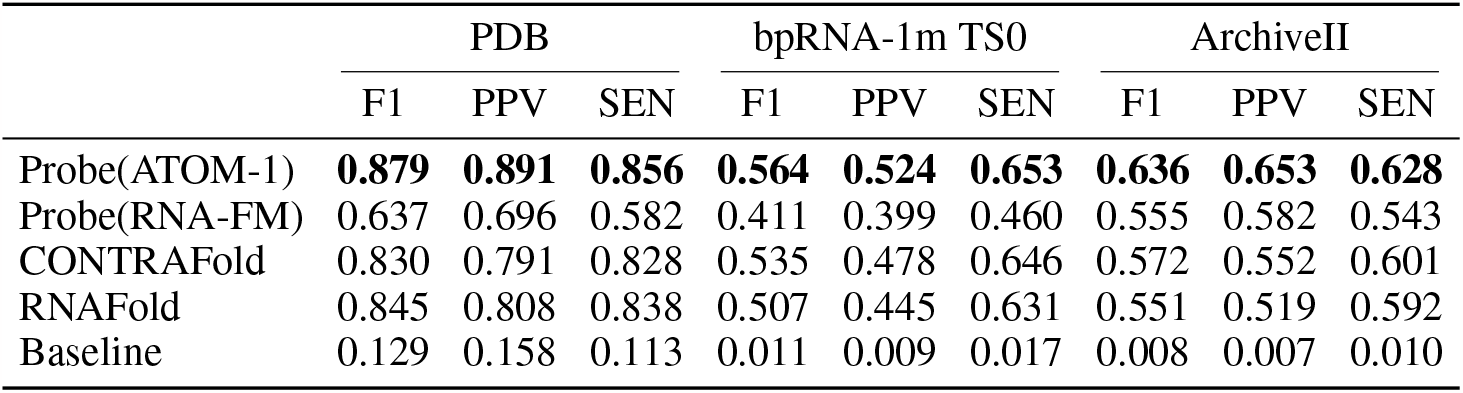
Our probe network applied to ATOM-1 achieves top accuracy on three secondary structure datasets. The table reports the mean of F1, PPV, and SEN applied to each of the three secondary structure test sets (PDB, bpRNA-1m TS0, Archive II) discussed in the main text.

|  | PDB |  |  | bpRNA-1m TS0 |  |  | ArchiveII |  |  |
| --- | --- | --- | --- | --- | --- | --- | --- | --- | --- |
|  | F1 | PPV | SEN | F1 | PPV | SEN | F1 | PPV | SEN |
| Probe(ATOM-1) | **0.879** | **0.891** | **0.856** | **0.564** | **0.524** | **0.653** | **0.636** | **0.653** | **0.628** |
| Probe(RNA-FM) | 0.637 | 0.696 | 0.582 | 0.411 | 0.399 | 0.460 | 0.555 | 0.582 | 0.543 |
| CONTRAFold | 0.830 | 0.791 | 0.828 | 0.535 | 0.478 | 0.646 | 0.572 | 0.552 | 0.601 |
| RNAFold | 0.845 | 0.808 | 0.838 | 0.507 | 0.445 | 0.631 | 0.551 | 0.519 | 0.592 |
| Baseline | 0.129 | 0.158 | 0.113 | 0.011 | 0.009 | 0.017 | 0.008 | 0.007 | 0.010 |

- Our PDB test set is constructed from single-chain unmodified RNA PDB entries deposited after May 1, 2020, resulting in 113 structures in 57 clusters.
- The ArchiveII test set was constructed by filtering out sequences longer than 1000 nucleotides, removing sixteen entries, with 3825 structures assigned to 1318 clusters.
- The bpRNA-1m TS0 test set was taken directly from https://zenodo.org/records/4430150, and after filtering had 1277 sequences assigned to 1276 clusters.

See the Supplementary Data for a reproducible list of all training PDB entries and test clusters.

### S2 3D structure prediction

#### S2.1 A simple 3D structure module

Inspired by Equifold [54], we model each RNA nucleotide as a collection of partially-overlapping rigid bodies, each represented as an element of **SE**(**3**). Because each RNA nucleotide has many degrees of freedom we use 8 rigid bodies for each nucleotide, though other, more parsimonious representations are possible. One peculiarity of 3D structure prediction is that this representation is not unique: an RNA structure represented by the series of rigid bodies *F*_1_, …, *F*_*n*_ *∈* **SE**(**3**) should be considered identical to *TF*_1_, …, *TF*_*n*_ for any *T ∈* **SE**(**3**) as this is a rigid transformation of the original structure.

This representation allows us to use a simple, parameter-free fixed nonlinearity to map from invariant vectors to 3D structures inspired by the Chroma structure module [55]. For every pair of rigid bodies *A* and *B* we predict the relative position of the origin of *B* in the coordinate frame of *A*: (*A*^*−*1^*B*)**0**. Here **0** is the zero vector in **R**^3^, and we use the natural action of **SE**(**3**) on **R**^3^. This vector (which we call the *pairwise displacement*) is invariant with respect to left action of **SE**(**3**) on itself by multiplication, as (*TA*)^*−*1^(*TB*) = *A*^*−*1^*B* for all *T ∈* **SE**(**3**), and is thus a sensible thing to predict.

To find the single structure most consistent with the predicted pairwise displacements **d**_*ij*_ *∈* **R**^3^ we consider the following optimization problem:

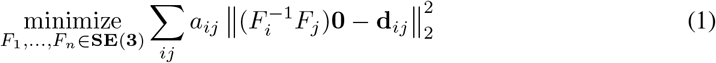

Here *a*_*ij*_ is a non-negative weight produced by our network.

While the optimization problem (1) does not have a closed-form solution it does allow for a very fast coordinate descent algorithm. This is because minimizing over a single rigid body *F*_*i*_ with all other bodies fixed is an instance of the celebrated Kabsch problem, which *does* admit a closed-form solution. To speed up convergence we use parallel block coordinate descent: we solve the Kabsch problem for each frame in parallel and update all frames simultaneously. During training or inference we run this algorithm until convergence or a maximum number of iterations (100) is reached. While parallel coordinate descent is not guaranteed to converge to the global optimum, we find that this procedure works in practice. To speed up and stabilize training we use a simple approximation to the derivative of this operation: we stop derivatives for all but the last step of the optimization procedure [56].

#### S2.2 Probe architecture

ATOM-1 embeds an RNA sequence of length *n* as a tuple of two arrays (*S, P*) with *S ∈* **R**^*n,d*^ and *P ∈* **R**^*n,n,d*^ ; following AlphaFold (AF2) [25], we refer to these as the single and pair representations respectively. The structure prediction probe takes these arrays as input features and applies three simple components: an input adapter, a small trunk, and the very simple structure module described above.

The input adapter is two linear layers that are applied independently along the last dimension of the single and pair representations to change *d* and *d*^*′*^. The trunk is an extremely shallow version of the Evoformer from AF2 (without any MSA features). In all of our experiments we use a trunk with two layers. Finally, the structure module applies linear mappings to the final dimension of the pair representations to produce the estimated displacements **d**_*ij*_ *∈* **R**^3^ and weights *a*_*ij*_, which are fed into the optimization problem 1 to produce a structure.

There are two additional input features used in some experiments: a sequence featurizer which embeds simple features of the input sequence (nucleotide identities and relative positions), and a recycling embedder, which is simply a pair of layernorms that can be used to feed the output of the main trunk as input to the trunk. All feature embeddings (the adapted ATOM-1 embeddings, sequence embeddings, and recycling embeddings) are summed before being fed into the trunk.

#### S2.3 Loss functions and training

We use a simplified version of the FAPE loss from AlphaFold [25]:

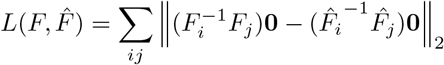

where *F* is the ground truth structure and 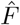 is the prediction. As is common in structure prediction we attach additional auxillary losses: we add a classifier head to the pair representation to predict the direction of the vector 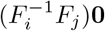 using a discretization of the sphere and an additional direct loss on the predicted displacement 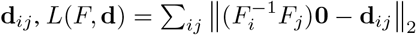

Our structure probe uses *recycling* [25]: during training we run the network a random number of iterations (between 0 and 3) without tracking gradients by feeding the output of the trunk into the network as an additional input. We then take one final step where we apply loss functions and compute the gradient with respect to the network parameters. During inference we run 10 recycling iterations before producing a structure.

#### S2.4 PDB split

We curate a set of PDB structures and a train/test split designed to make fair comparisons with competing 3D structure prediction models. We compile a structure set from all RNA-containing PDB structures, separating each PDB into groups of interacting chains and excluding cases where we expect RNA structure to be primarily determined by protein or DNA binding-partners. We then create a training set by selecting structures from PDB entries published prior to April 30, 2020. Since competitor methods may have been trained on structures released after April 30, 2020, we use only PDBs published after May 1, 2022 as test structures. We finally apply a sequence similarity filter (<80% similarity as measured by `cd-hit-est-2d` [57]) to ensure that our test set is sufficiently far from our training set.

We report results on clusters of test set sequences, grouping sequences with similarity >80% as measured by `mmseqs2` [58].

A list of PDB IDs in our train and test set is available in the Supplementary Data.

### S3 In-solution stability

#### S3.1 Dataset

In 2020 the Das lab at Stanford launched the OpenVaccine Kaggle challenge to accelerate the development of computational tools for vaccine design [50]. Participants were asked to predict nucleotide-level degradation rates in either high temperature or high pH, with and without high concentrations of magnesium, as well as reactivity when exposed to a small-molecule reagent.

The OpenVaccine organizers provided experimental labels for 2400 107-nt-long constructs as the training set. Upon completion of the challenge, participating teams were evaluated based on the accuracy of their predictions on a test set of 1172 130-nucleotide-long sequences. These sequences comprised a *private test set* during the challenge *i*.*e*., participants did not have access to individual labels or aggregate prediction accuracy on this set during the challenge. Instead, participants calibrated their methods based on an aggregate accuracy score for 629 107-nucleotide-long sequences in a *public test set*. In the main text we refer to these public and private test sets as the *validation* and *test* sets respectively, due their roles in the competition. Further details on the datasets, as well as links to the data itself can be found in the follow-up paper from the challenge organizers [51].

#### S3.2 Prediction task

Given a sequence, the prediction target for this task is a simultaneous, per-nucleotide estimate of three different experimental measurements. Predictions were only scored up to a certain number of nucleotides, which we denote here as *L* (68 for the training set, 91 for the private test set). Given a prediction as well as a corresponding set of experimental measurements *ŷ, y ∈* **R**^*L×*3^, the per-sample, mean columnwise root mean squared error (MCRMSE) is computed as

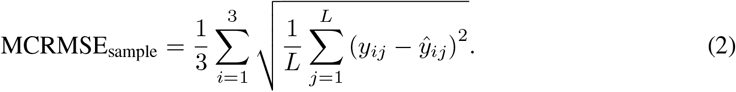

In the challenge, the aggregate MCRMSE over the dataset was computed in the following way:

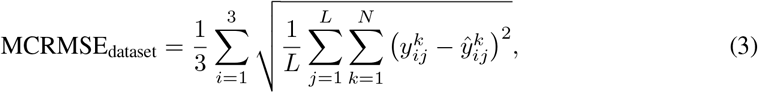

where *N* is the number of samples in the dataset, and *y*^*k*^ denotes the value of the *i*^th^ condition at position *j* for sample *k*.

#### S3.3 Probe architecture

Following the previous sections, our probe architecture takes in single and pair representations from ATOM-1 as input, and consists of an input adaptor, trunk, and prediction head. The input adaptor of the probe consists of linear layers applied to the sequence dimensions of the single and pair representations. The trunk of the probe uses a multilayer perceptron (MLP) to compute attention weights for the pair representation. These attention weights are used to compute a multi-head attention residual update to the single representation. The single representation then receives a residual update from a one-layer MLP. Finally, a linear layer projects the updated single representation down to the output dimensions to generate the predictions. In total, the prediction model contains about 10 million parameters.

### S4 External network evaluation

We make predictions with two external 3D structure models: RhoFold [38] (formerly known as E2EFold-3D), and RoseTTAFold2NA [39], and one external foundation model RNA-FM [22]. In this subsection we briefly describe how we used these models.

For RoseTTAFold2NA, we used commit `43bdd89`, dated November 9, 2022, from the GitHub repository (https://github.com/uw-ipd/RoseTTAFold2NA/tree/main) and weights `RF2NA_sep22.pt`. We prepared all conda environments and MSA databases as described in the README file. To make predictions we use the prediction script provided, which automatically generates MSAs.

For RhoFold, the original GitHub repository https://github.com/ml4bio/E2Efold-3D was moved, but the new repository was not available at the time of writing. Instead we used a cached copy of the code and weights from October 10, 2022, and exactly reproduced the conda environment as described in the README. We re-used the MSAs generated as part of the RoseTTAFold2NA workflow.

For RNA-FM, we used commit `3e24749` of the GitHub repository https://github.com/ml4bio/RNA-FM. The weights were downloaded prior to August 24, 2022. For inference we used the pretrained network configuration `extract_embedding.yml` and the corresponding weight file `RNA-FM_pretrained.pth` to avoid data leakage with the secondary structure test sets considered. To extract embeddings, we directly called the model and extracted the array corresponding to the key `representations` from the `results` dictionary. In order to extract the attention weights we toggled the `flag need_head_weights`, and used the value corresponding to the `attentions key`. We confirmed that the single representation embeddings used were correct by comparing to the RNA-FM web server.

