## Supplementary material for "ATOM-1: A Foundation Model for RNA Structure and Function Built on Chemical Mapping Data": README.txt

# pdb_2d_split.csv
PDB secondary structure split and data file

Column descriptions:

"pdb_id": PDB entry the sequence originated from. This is not unique for the train set, as some PDB entries may have up to four secondary structures in the train test.
"train_or_test": Specifies if the example is in the train set or test set.
"test_cluster_idx": The index of the cluster for the test set. This is not applicable for the train set.
"sequence": The RNA sequence.
"structure": The RNA secondary structure in dot-bracket notation.

# rfam_2d_split.csv
Secondary structure test data from the Rfam-based datasets bpRNA-1m and ArchiveII.
Base pairs were extracted from the bpseq files located at in the data file `mxfold2_data.zip` found at https://zenodo.org/records/4430150. 

Column descriptions:

"name": The name of the RNA sequence.
"dataset": Specifies if the example is in the train set or test set.
"test_cluster_idx": The index of the cluster for the test set. These are duplicated between the two datasets, so the datasets must be separated before clustering metrics.
"bpseq_path": The path to the bpseq file in the `mxfold2_data.zip` file.

# pdb_3d_split.csv
PDB secondary structure test data from the RNA-Puzzles 13 dataset.

Column descriptions:

"pdb_id": PDB entry the sequence originated from. This is a unique list of PDBs for the test set without curation of which chains were used.
"train_or_test": Specifies if the example is in the train set or test set.
"test_cluster_idx": The index of the cluster for the test set. This is not applicable for the train set.
